# Probabilistic estimation of short sequence expression using RNA-Seq data and the “positional bootstrap”

**DOI:** 10.1101/046474

**Authors:** Hui Y. Xiong, Leo J. Lee, Hannes Bretschneider, Jiexin Gao, Nebojsa Jojic, Brendan J. Frey

**Affiliations:** Department of Electrical and Computer Engineering, University of Toronto, Toronto, Ontario M5S 3G4; Banting and Best Department of Medical Research, University of Toronto, Toronto, Ontario M5S 3E1; Microsoft Research, Redmond, WA 98052, USA; Canadian Institute for Advanced Research, Toronto, Ontario, M5G 1Z8, Canada

## Abstract

When estimating expression of a transcript or part of a transcript using RNA-seq data, it is commonly assumed that reads are generated uniformly from positions within the transcript. While this assumption is acceptable for long transcript sequences where reads from many positions are averaged, it frequently leads to large errors for short sequences, *e.g*., less than 100 bp. Analysis of short sequences, such as when studying splice junctions and microRNAs, is increasingly important and necessitates addressing errors in short-sequence expression estimation. Indeed, when we examined RNA-seq data from diverse studies, we found that large errors are introduced by variations in RNA-seq coverage due to sequence content, experimental conditions and sample preparation.

We developed a technique that we call the positional bootstrap, which quantifies the level of uncertainty in expression induced by non-uniform coverage. Unlike methods that attempt to correct for biases in coverage, but do so by making strong assumptions about the form of those biases, the positional bootstrap can quantify the noise induced by all types of bias, including unknown ones. Results obtained using independently generated RNA-seq datasets show that the positional bootstrap increases the accuracy of estimates of alternative splicing levels, tissue-differential alternative splicing and tissue differential expression, by a factor of up to 10.

A Python implementation of the algorithm to quantify splicing levels is freely available from github.com/PSI-Lab/BENTO-Seq.

## 1 Introduction

We describe a simple, novel procedure that can significantly increase the usefulness of data generated using massively parallel RNA-seq technologies. In support of downstream research and biotechnologies, several computational techniques have been developed so as to estimate the abundances, or relative abundances, of short transcript sequences using RNA-seq data (Jiang and Wong, 2009; Li *et al*., 2010a; Li and Dewey, 2011; Trapnell *et al*., 2010; Roberts *et al*., 2011; Huang *et al*., 2013; Jiang and Salzman, 2013; Hu *et al*., 2013). Despite extensive research and technology development, this remains a challenging problem because: complete libraries of transcripts are not available, even for cell types of major interest such as normal human cells and cancer cells; short read lengths and fragment sizes often make it impossible to uniquely resolve transcripts, leading to errors (Lacroix *et al*., 2008; Steijger *et al*., 2013); technical artifacts introduced by sequencing protocols are not well understood, cannot be properly modelled, and introduce significant noise, which is further compounded by the two above-mentioned issues (Dohm *et al*., 2008; Hansen *et al*., 2010; Zheng *et al*., 2011); and there are no widely accepted, gold-standard datasets that can be used to convincingly benchmark techniques for estimating expression. Improvements in protocols have partly addressed some of these challenges, such as uniformity of read coverage (Mortazavi *et al*., 2008; Levin *et al*., 2010), but computational methods are indispensable for improving estimation accuracy (Li *et al*., 2010b; Hansen *et al*., 2010; Roberts *et al*., 2011; Li and Dewey, 2011; Jones *et al*., 2012; Jiang and Salzman, 2013; Hu *et al*., 2013).

Because of the practical need for abundance estimation, the above challenges are usually dealt with computationally by using data models that are known to be inaccurate, or by ignoring the problems altogether. Noise models may partly address artifacts introduced by sample preparation, hexamer priming, PCR amplification, RNA secondary structure, relative distance to the 3’ end of the transcript, local sequence composition, such as GC-content, and biases from the mapping stage (Dohm *et al*., 2008; Hansen *et al*., 2010; Zheng *et al*., 2011). However, inaccuracies in data models could also make matters worse. Methods that supposedly correct the ‘noise’ introduced by local sequence composition are developed and applied, even though this approach corrupts the underlying expression signal, which is itself, of course, a function of the transcript sequence. Computational methods are routinely compared using simulated data, even though it is widely known that such data does not accurately reflect real data. Consequently, expression is often estimated by simply summing up the reads that uniquely map to a transcript.

Here, we take a different approach. Instead of trying to correct errors using inaccurate data models, we introduce the ‘positional bootstrap’, which quantifies the uncertainties in abundance estimates in a way that works well for many different sources of noise, including unknown ones. This enables downstream analyses to properly prioritize or weight abundance estimates. For example, given an RNA-seq dataset and a set of transcript sequences, our method can be used to rank the abundance estimates. Below, we demonstrate the severity of different types of noise using reads mapped to exon-exon junctions, describe the positional bootstrap, and then examine the application of our method to estimating alternative splicing levels and junction expression levels. We conclude with a brief discussion about the general applicability of our method to other estimation tasks as well as how it can be combined with existing bias-correction methods.

## 2 Biases in RNA-seq

To illustrate the effects of noise introduced by various biases, including across different sample preparations, we analyzed the 75bp Illumina Bodymap dataset and the 76bp human dataset from the Kaessmann laboratory (Brawand *et al*., 2011). These datasets were derived from poly-A selected mRNA of healthy individuals and have four overlapping tissue types: brain, heart, kidney and liver. We mapped the two datasets to 671,448 exon-exon junctions derived from hg19 annotations, using TopHat with the same parameter settings. We required each junction read to overlap at least 8bp with both exons, making the number of possible read mapping positions for each junction to be 60 for Bodymap and 61 for Kaessmann’s dataset. For a given junction, if there were no noise sources that lead to positional variations in read coverage, the underlying transcript abundance would be the same across different positions, as would the Poisson parameter that is assumed to generate the reads. However, as described below, we observed widespread over-dispersion in the number of reads mapped to junction positions, indicating the existence of considerable position-dependent noise.

To examine the strength of different biases, for each exon-exon junction and tissue in the Bodymap data, we counted the number of unique positions with reads, *m, i.e*., positions that had one or more mapped reads. It is interesting to examine this quantity because, in contrast to the total number of reads that map to a junction, it is much less sensitive to ‘read stacks’ and other biases. For example, a read stack is only counted once in *m*. Comparing the observed distribution of *m* to a simulated distribution assuming uniform coverage can reveal the strength of sequencing biases. For each junction and tissue, we used the actual number of mapped reads, to simulate bias-free data by assuming that reads map to all positions in an unbiased, or uniform, manner (see SM). Fig. 1A shows the distribution of the number of unique positions, *m*, for the actual data (red), compared to the bias-free, simulated data (blue), for all junctions where the total number of mapped reads is between 10 and 20. The two distributions are very different, indicating that the approach of summing up the reads over positions is highly flawed.

A common misconception in the community is that if the number of mapped reads is larger, the relative noise level is lower. To explore this, in Fig. 1B we plot the distribution of *m* for the actual and bias-free data, when the total number of mapped reads is between 50 and 60. We see that the distributions overlap even less for these high-count cases.

We additionally found that the actual data contains both sequence- and experiment-dependent biases. This is observed by first mixing the experiments to cancel out the majority of experiment-dependent bias, and then perform the above analysis of comparing the observed distribution of *m* (green lines in Fig. 1 A-B) to the distribution assuming bias-free read mapping. For every experiment and every short sequence, we sampled *n* reads that are mapped to the event from all 16 experiments. The resulting data still contains sequence-dependent biases that are shared across experiments, because we only sampled from actual, mapped reads. For 10 ≤ *n* ≤ 20 in Fig. 1A, this distribution (green) is located roughly 2/3 between the distribution for data containing both sequence- and experiment-dependent biases and the distribution for bias-free data. Because all experiments are measuring the same sequences and the same RNA species, this suggests that about 2/3 of the overall bias is experiment-dependent and the other 1/3 is sequence-dependent. For 50 ≤ *n* ≤ 60, the distribution of *m* is plotted in Fig. 1B (green), and indicates that when the number of mapped reads is higher, sequence dependent bias plays more of a role.

**Figure 1.**
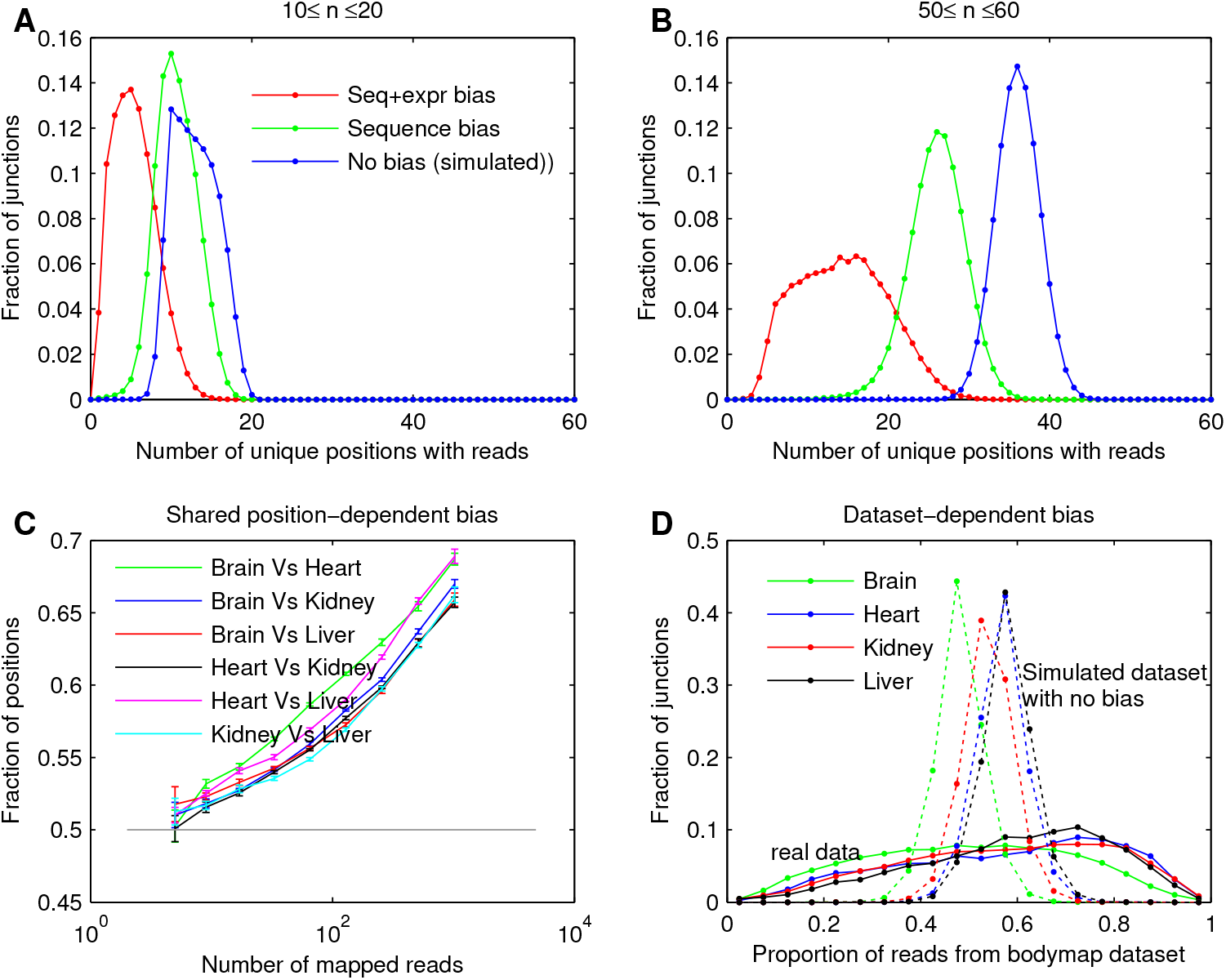
Sequence-, experiment-, and dataset-dependent biases in real RNA-seq data. A-B: Distribution of the number of positions with at least one mapped read spanning an exon-exon junction in the Bodymap data (red), randomly resampled Bodymap data with effects of experiment-dependent bias canceled (green), and simulated data with no sequence- and experiment-dependent bias (blue). *n* is the total number of reads mapped to the junction. C: For each pair of tissues, the observed frequency of positions in pairs of tissues that have more reads than the median. All lines are above 0.5, showing reads tend to be mapped to the same positions in two tissues. D: Distribution of junctions by the proportion of reads coming from the Bodymap dataset versus Kaessman’s dataset. Solid line: RNA-seq data. Dotted line: expected distribution if there exists no dataset-dependent differences across junctions. The real data have a much wider distribution, reflecting dataset-dependent differences in sequencing technology and sample variability.

To further show the effects of sequence bias, we examined whether reads tend to be mapped to the same positions in two different tissues, *i.e*., whether different tissues have a shared sequence-dependent bias. Fig. 1C plots the observed frequency of positions in pairs of tissues that have more reads than the median. The frequency exceeds what would be expected at random (0.5), especially for larger *n*, when the effects of sequence bias are expected to dominate, as shown above.

Finally, to explore dataset-dependent biases, for each junction, we compared the number of reads mapped from the Bodymap and Kaessman datasets and computed the proportion of reads from the Bodymap dataset. This quantity was computed for both the actual and simulated bias-free data, and the distributions are shown in Fig. 1D (solid lines for real data and dashed lines for simulated data). The distributions dervied from the actual data have a much wider variance than those from the simulated bias-free data, indicating that there are significant dataset-dependent biases.

## 3 The positional bootstrap

Our technique, the positional bootstrap, is very simple to implement and quickly assesses uncertainty in transcript abundance estimates induced by position-dependent or sequence-dependent biases. It is based on the bootstrap, a highly robust non-parametric method used for estimating uncertainty. In the following, we briefly review the bootstrap procedure and apply it to the estimation of short sequence abundances and splicing levels.

For a given transcript sequence, let *R* be the set of potential positions where corresponding short reads can be mapped, let 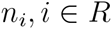 be the number of reads that map to position *i* in the RNA-seq dataset, and let 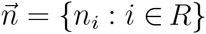. In this paper, we refer to the ‘mapped position’ of a read as the position of the first base of the RNA-seq read so that all reads mapped to a certain position are the same. As a result, reads mapped to the same position share the same sequencing bias and *n_i_* is affected by these biases differently.

Suppose there is a quantity such as expression, λ, that we are interested in estimating and we have a technique that takes the dataset 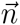 as input and outputs an estimate 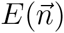 of λ. For example, λ might be the transcript abundance and the estimate might be the average number of mapped reads 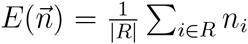 or the median number of mapped reads across the positions. For a true parameter value λ and a particular sequencing procedure, there is a distribution of possible datasets that we may obtain. If we apply the estimator to these datasets, we obtain a probabilistic distribution of the estimate 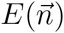. By examining how close the distribution of 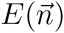 is to the true parameter value λ, we can draw conclusions about how well the experimental procedure combined with the computational technique estimates λ. In particular, from this distribution, we can calculate the estimator variance and construct confidence intervals.

To obtain an estimate of the quality of 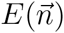 using the above procedure, we have to repeat the experiment a large number of times to obtain a distribution of 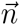 and 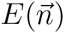. Practically, this approach is infeasible since all the data samples are usually combined into a single dataset to produce the best estimate. As a result, there is only one dataset and one estimate whose quality we seek to determine. The Bootstrap is an ingenious method proposed by Efron (Efron, 1982) to approximately generate new datasets from a single original dataset. This is accomplished by using the empirical distribution of data points in the collected dataset as an approximation to the population distribution of all possible data points. From this empirical distribution, data points are re-sampled with replacement to create new datasets. Using these re-sampled datasets, a distribution of estimators is produced. Because there is usually a considerable amount of independently collected data points, the empirical distribution approximates true population reasonably well, which is the only assumption made in the bootstrap procedure. In this situation, the datasets generated by bootstrap are good approximations of hypothetical new datasets. This in turn makes the computed distribution of estimators close to the true distribution. In many applications, the bootstrap has been very successful because of its robustness and ease of use.

In our application, the number of reads (*n_i_*) mapped to positions corresponding to a certain short sequence are treated as data points. These positions have mapped reads that originated from the same RNA species with a certain abundance but may be affected by different sequencing artifacts. Therefore, *n_i_* comes from a certain distribution with a shared mean corresponding to expression and variations corresponding to sequencing bias and sampling variability. We bootstrap these positions to form sample datasets with biases that are typically encountered during sequencing.

Each bootstrap dataset is generated by re-sampling the positions in *R* with replacement while keeping the total number of sampled positions the same as in the original dataset. An illustration of this procedure is shown in the first two rows of Fig. 2. We denote the index of the *i*-th data point in the *k*-th bootstrap dataset sample as *I*^(*k*)^(*i*) so that the entire *k*-th bootstrap dataset is 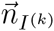. The *i*-th element in *I*^(*k*)^ is generated as an independent random sample from *R:*

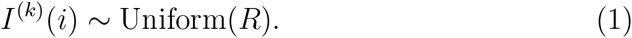

To obtain good bootstrap estimates, the number of bootstrap samples, *K*, should be large to cover many likely datasets. After bootstraping the datasets, each dataset is processed to produce a point estimate 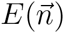 for λ or a Bayesian posterior distribution 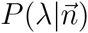 for λ. During this estimation procedure, read generation is assumed to be bias-free and follow standard distributions such as the Poisson or binomial distributions. As a result, the bootstrap procedure effectively handles the variation due to sequencing bias and the estimation procedure handles the inference assuming a bias-free read generation process.

If point estimates are used, the standard deviation across the bootstrap samples can be used as a confidence measure. If a Bayesian inference procedure is used for λ, each bootstrap sample produces a distribution over λ. In this case, we proposed a ‘posterior distribution with bootstrap’, 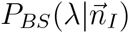, which combines the posterior probability of all of the bootstrap samples:

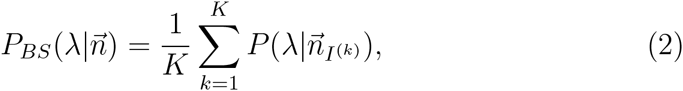

where *n_I_*(*k*) is the *k*-th bootstrap dataset. To obtain a confidence measure, we can use the variance of λ in 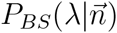, which is equal to the average variance of λ in 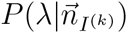 plus the variance of the expected value of λ across the bootstrap samples.

**Figure 2.**
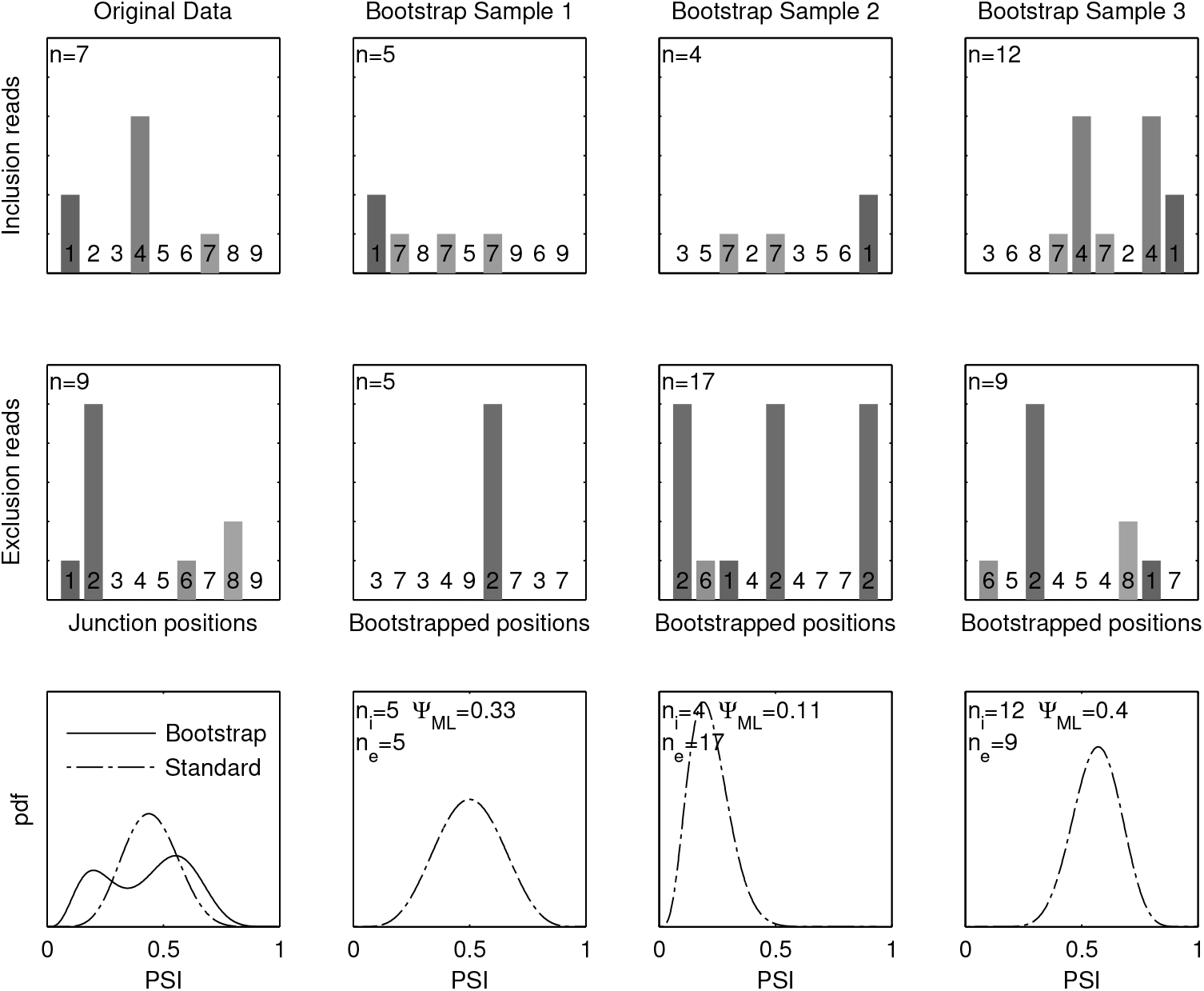
An illustration of the positional bootstrap. First two rows each represent a junction with nine mappable positions and the number of reads mapped to them. The left column is the original data and the rest are the bootstrap samples. In each bootstrap sample, the positions (shown above the x-axis) in the original data is sampled with replacement, such that repeats and skips almost always occur. Each bootstrap sample is processed to obtain a distribution of PSI values shown in the bottom row with the standard method. The distributions from bootstrap samples are averaged to produce the final bootstrap distribution of PSI, shown as the solid curve in the bottom left panel in comparison with the distribution produced by standard method with the original data (broken curve).

## 4 Estimators for expression and alternative splicing

Here, we show how the above technique can be used to obtain distributions of estimators for transcript abundance and relative isoform levels for alternatively spliced transcripts.

### 4.1 Expression estimation

Following the notation above, the number of reads mapped to a set of positions is {*n_i_*}. Assuming there is no sequencing bias, and that the number of reads is generated from a Poisson distribution with underlying abundance λ, the likelihood function of the entire dataset is:

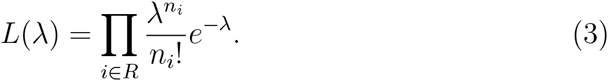

Let 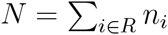 be the total number of reads and *P = |R|* be the total number of positions considered. We can use the maximum likelihood estimator as a point estimator, which is 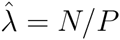. In addition, we can use a Bayesian framework with an improper uniform prior over λ. Then the unnormalized posterior distribution is the same as the likelihood function. Rearranging it, we obtain:

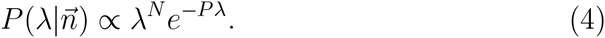

From the above, we observe that the posterior is the Gamma distribution with a shape parameter equal to *N +* 1 and a scale parameter equal to 1/*P*. The variance is equal to (*N +* 1)/*P*^2^, which can be used as the confidence for this estimator.

### 4.2 Splicing level estimation

In this section, we propose a Bayesian framework for the estimation of relative abundances of two splicing isoforms. We consider two spliced RNA products that are locally different. For example, in cassette splicing, the middle exon B in a triplet of exons A-B-C might be skipped, resulting in either the inclusion variant A-B-C or the exclusion variant A-C. For these two variants, we first identify two sets of positions to which mapped RNA short reads can only originate from one of the variants. We refer to these two sets of positions as the ‘inclusion positions’ and the ‘exclusion positions’. In the cassette splicing example, the inclusion positions corresponds to reads that span the A-B junction, the B-C junction and reads mapped to the body of B, while exclusion positions correspond to reads that span the A-C junction.

We only use the numbers of reads mapped to the inclusion and exclusion positions defined above so there is no uncertainty about which isoform a read came from. If a position is found to be not mappable, possibly due to reasons such as sequence repeats, the position can be discarded. In addition, positions whose mapped reads do not span an exon-exon junction can be discarded to avoid error introduced by unspliced reads. These procedures reduce the set of inclusion and exclusion positions, but do not change the inference procedure presented here. In the following analysis, the input data consists of two sets of counts corresponding to the number of reads mapped to the inclusion and exclusion positions. The size of these two sets of positions need not to be the same and the goal is to determine the distribution over the percent of transcripts with the central piece spliced in (PSI), as generally defined for different alternative splicing events in (Wang *et al*, 2008).

Let *I* and *E* be the set of inclusion and exclusion positions and the number of reads mapped to position *i* be *n_i_*. Suppose in the biological sample, the combined expression of the inclusion and exclusion variant is *β* and the PSI value is *ψ*. Then, by the definition of PSI, the abundance of the inclusion variant is *ψβ* and the abundance of the exclusion variant is (1*-ψ*)*β*. Based on the assumption that short RNA-seq reads are independently generated with a Poisson parameter proportional to the abundance, the combined likelihood of the observed RNA-seq read data is:

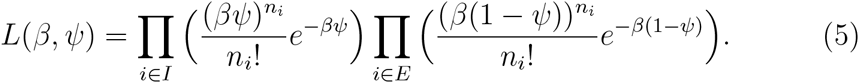

Rearranging the above equation, we obtain:

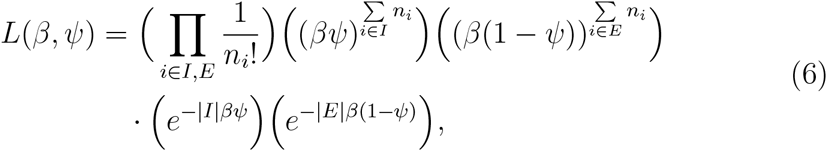

where |*I*| is the size of *I* and |*E*| is the size of *E*. Clearly, the likelihood function depends on 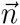 only through the total number of inclusion and exclusion reads. To simplify the notation, let 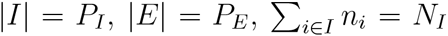 and 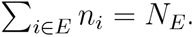. For each event, these four counts are the sufficient statistics that jointly determine the likelihood function of *β* and *ψ*. Using this notation and rearranging, the likelihood function is:

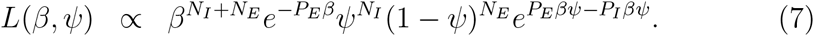

Note that when the number of the inclusion positions is the same as the number of exclusion positions (*P_I_* = *P_E_*), the last term is constant and the likelihood function is separable between *β* and *ψ*. The part depending on *ψ* is a standard binomial likelihood function. Practically, this can happen if we only consider one junction for each variant and all junction positions are mappable. Intuitively, this is because when either an inclusion or exclusion read is observed, the probability that it is an inclusion read is the same as PSI without scaling by the number of possible inclusion/exclusion positions.

However, there is dependency between *ψ* and *β* in general because of unequal number of positions. To obtain the posterior distribution of *ψ*, we need to integrate out *β* from the joint posterior distribution of *β* and *ψ*. Here, we choose the Beta prior with parameters *a* and *b* for *ψ* and an independent exponential prior with parameter *r* for *β:*

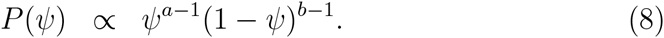

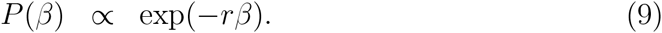

The two parameters *a* and *b* in the Beta prior represent how likely the inclusion variant appears in general and how strong the prior is. They can be inferred from data using an empirical Bayes approach. Furthermore, different Beta priors can be used for different events, if additional information is available. Similarly, the parameter *r* in the exponential prior for expression represents a likely expression level of a short sequence scaled by sequencing coverage. Without specific prior knowledge, we can use the uniform prior as a the default setting with *a = b =* 1 and *r* = 0. Note that when *r* = 0, the prior of expression is improper as it cannot be normalized, but with observed data, it becomes normalizable. Using these priors, the unnormalized posterior distribution is:

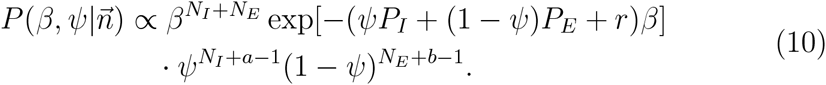

We integrate over *β* to obtain the posterior distribution of *ψ*. As long as *r* ≥ 0, the integral over *β* converges when there is at least one mappable inclusion or exclusion position.

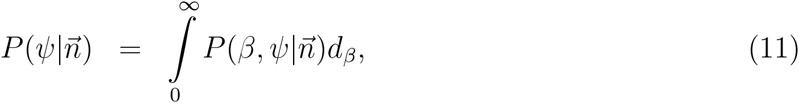

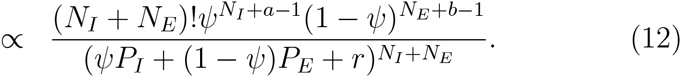

The above posterior distribution is a rational function of degree equal to the total number of mapped reads, which can be large for highly expressed genes. To obtain exact samples or exact posterior distribution would require integrating ψ to obtain the normalizing constant of *P*(*ψ|*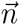). This can be slow if many reads are mapped. To speed up computation, we observe that *ψ* is one dimensional and bounded between 0 and 1. This makes the distribution of *ψ* convenient to approximate using a grid. We use a uniform grid with linear interpolation to approximate the probability density function of the posterior. This way, the distribution of *ψ* can be obtained in constant computation time with respect to *N* and *P*. To prevent numerical underflow, the standard log-sum-exp technique is used. The number of points in the grid can be set by the user, and a default value of 500 is used in our software implementation.

## 5 Results

To demonstrate the effectiveness of the proposed bootstrap method, we used it to estimate PSI for cassette splicing, which is the most common type of alternative splicing. In cassette splicing, the alternative exon B in an consecutive exon triplet A-B-C can be excluded while exons A and B are assumed to be constitutive. We mapped the Bodymap and the Kaessmann datasets described above to 43,848 cassette exon triplets and only used the junction reads. The bootstrap method was applied with the uniform prior on both PSI and expression (*a* = *b* = 1, *r* = 0). In addition, we applied Bayesian PSI inference without the bootstrap, which is denoted as the ‘standard method’.

Fig. 3 compares the mean and standard deviation obtained using the bootstrap procedure and the standard method, and includes the details for two examples. In these two examples, total number of mapped reads are similar, but the reads mapped to one exon is much more concentrated on certain spots, suggesting sequencing bias. In the result, PSI mean is similar for the two methods, but the standard deviation is quite different, because sequencing artifacts are detected. Fig. 4 shows a scatter plot of the additional variance captured by the bootstrap method compared to the standard method for both datasets. We observe that there is a positive correlation due to a sequence-dependent bias that is shared between the two datasets. However, significant differences exist due to an experiment-dependent bias and sequence-dependent biases that are different between the two datasets.

**Figure 3.**
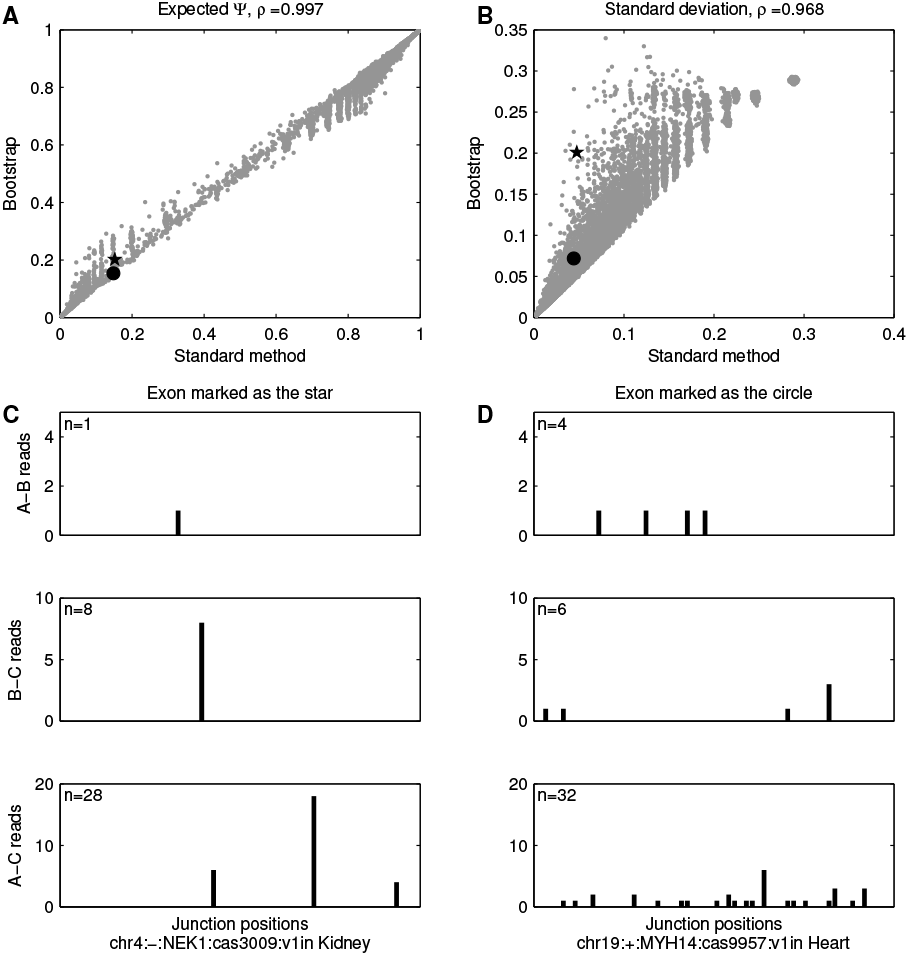
A-B: Comparison between the estimated PSI and its standard deviation obtained by the bootstrap method and the standard method. Pearson correlation coefficient is shown above the figures. Correlation is very high for expected PSI while bootstrap increases the standard deviation estimations for many exons due to unevenly mapped reads. C-D: Real RNA-seq read distributions of two example exons, shown as circle and star in A-B, having similar PSI values and standard deviation estimations from the standard method. However, the bootstrap accounts for the high unevenness of read mapping in C and assigns a much higher standard deviation.

**Figure 4.**
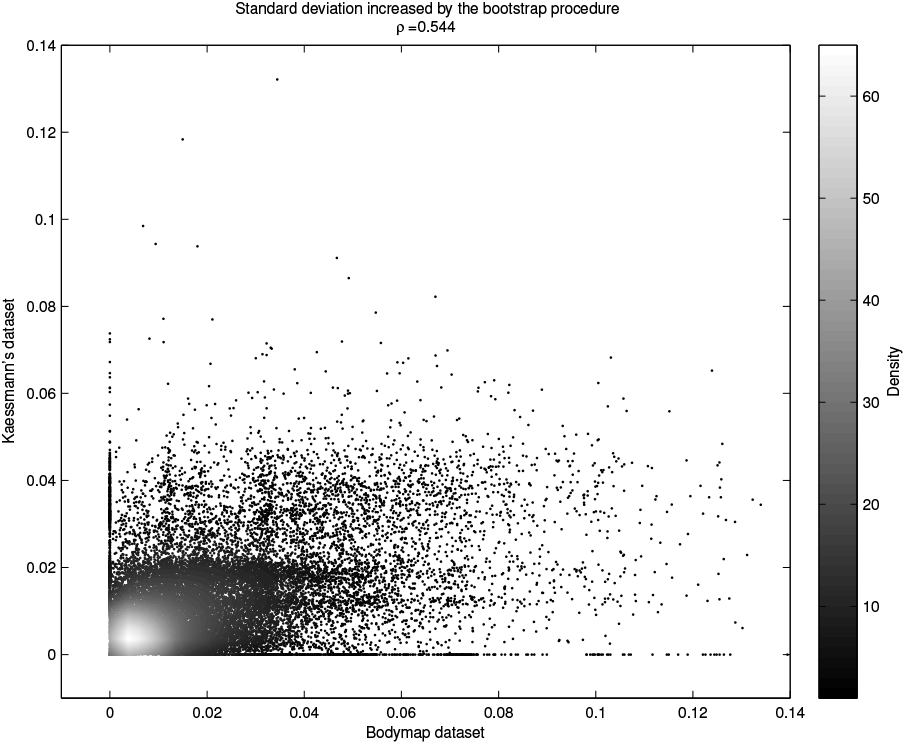
Additional PSI variance estimated by the bootstrap method compared to the standard method for two datasets. Points at the origin correspond to exon-tissue combinations whose bootstrap confidence estimate is the same as the standard confidence estimate. Points away from the origin correspond to exon-tissue combinations that have a higher bootstrap confidence estimate, because biases are accounted for.

We used several methods to evaluate the performance gain produced by the bootstrap procedure. First, we computed the correlation between PSI estimates from two datasets (Bodymap and Kaessmann) for the top *k* percent most confident exons, ranked by the confidence estimate produced by the standard method and the bootstrap method (Fig. 5 A). We observe the correlation of PSI decreases as more exons are included, indicating that both confidence estimates are representative of uncertainty. However, the bootstrap confidence is significantly better and decreases the error (1-correlation coefficient) by a factor of 2 for the top 10% of exons.

**Figure 5.**
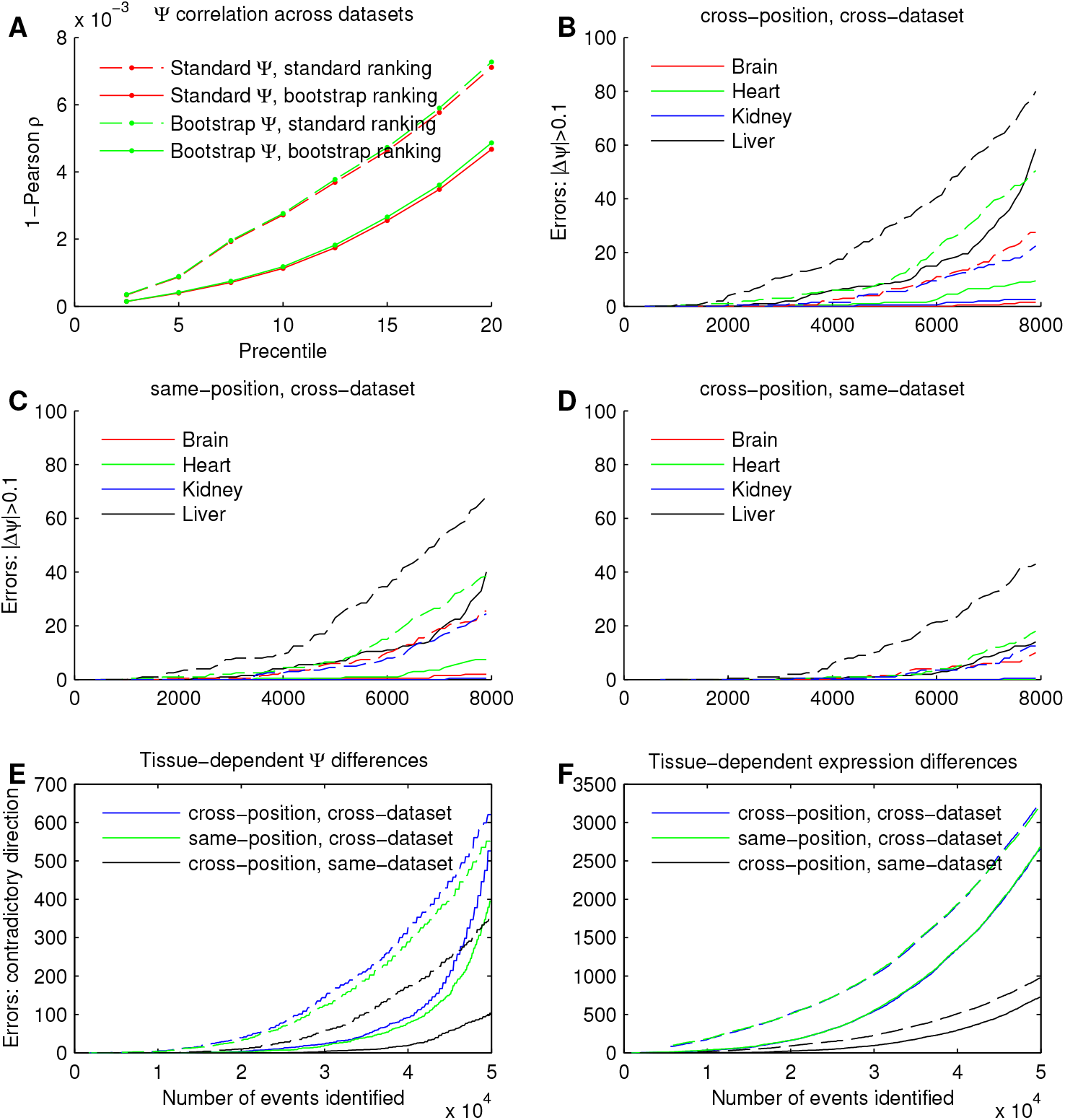
A. Cross-dataset Pearson correlation coefficient of the k percent most confident PSI estimates. The raking is computed by sorting the standard deviation in increasing order. Using the bootstrap standard deviation to rank events increased the correlation of PSI values estimated from two independently prepared datasets by both methods. The version of PSI estimate used made little difference. B-F: Number of disagreements (errors) of PSI and junction expression estimates when comparing different divisions of datasets. Dataset divisions are described in the main text and SI. Events are sorted by increasing standard deviation. Dotted line: standard method. Solid line: bootstrap method. B-D: Number of errors in PSI estimation. An error is defined as PSI difference greater than 10% between two datasets. E-F: error rates of tissue dependent PSI differences and junction expression differences. In A-F, using the bootstrap to estima1t6e confidence (solid line) appear to always produce lower error rates than the standard method, indicating that the bootstrap captured genuine inaccuracies produced by RNA-seq technology.

Next, we split each dataset into two halves by junction positions (positions 1-30 and 31-60) so that the positions covered by each half are different. These two halves represent two junction datasets that may be produced by a technology with a shorter read length, but with the same distribution of position-dependent sequencing biases as well as other bias introduced during upstream processing, such as during sample preparation and amplification. However, their idiosyncratic biases are different due to different sequences associated with different positions. Since these two halves are measuring the same RNA species, a robust procedure that properly takes into account uncertainty introduced by sequencing bias should produce little difference when estimating PSI. Similarly, PSI values estimated from the same half of the Bodymap and Kaessmann’s dataset and the PSI values estimated from different halves of the different datasets can be compared. For these two comparisons, genuine PSI differences might exist because of possible biological difference between the samples. However, dataset-dependent sequencing bias, whose severity is shown above in Fig. 2 D, can also produce apparent sample-dependent PSI differences that are erroneous. Therefore, a robust method that captures sequencing bias is expected to produce fewer confident differences in PSI.

Using these two halves of the two datasets, four sets of PSI values and confidence estimates were produced using the standard method and the bootstrap method. For each method, we analyzed the agreement of all six pairs of these four estimates by separating them into three categories based on if the position and the dataset are the same (Fig. 5 B-D). Exons are sorted in descending order of confidence in PSI and an error is defined by a greater than 10% disagreement in the PSI value. As expected, the cross-position, cross-dataset comparison (Fig 5 B) produces the largest difference, because both the experiment and the sequence are different. The same-position, cross-dataset comparison produced more error than then cross-position, same-dataset comparison, suggesting there is more experiment-dependent bias than sequence-dependent bias. For all of these experiments, compared to the standard method, the bootstrap method made significantly fewer errors in estimating PSI for the exons to which it assigned highest confidence.

We also evaluated the proposed bootstrap method in terms of detecting tissue-dependent PSI differences and tissue-dependent exon-exon junction expression differences. All pairs of tissues are considered, producing a total of 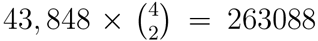 exon-tissue-pair combinations. Z-scores of the expected PSI differences are used to detect tissue specific splicing changes:

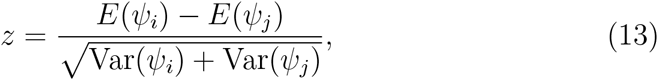

where *ψ_i_* is the PSI value estimated for the *i*-th tissue. These z-scores are computed for all exon-tissue-pair combinations and ranked by their absolute value. To detect a set of *k* most significant tissue-specific splicing changes, a threshold on the absolute value of z-scores corresponding to *k* is used. Using the setup described above, we used the same event-tissue-pair combination from two datasets such that they differ in position or origin to verify the detection of tissue-dependent splicing differences. The top *k* events are used for both datasets and an error is detected if the direction of change is different between the two datasets. Using this criteria, we compared the performance between the bootstrap method and the standard method. As shown in Fig. 5 E, the bootstrap method produced a much lower error rate for the most confident events than the standard method. In addition, the bootstrap method can be used to detect junction expression differences. For this task, the mean of the posterior distribution of log(*β*) is used as the expression and the variance of log(*β*) is used as the confidence estimate. Z-scores are used to rank the events. Again, a much higher detection rate of tissue-dependent difference is achieved by the bootstrap method.

## 6 Conclusions

Sequencing artifacts are prevalent in RNA-seq, they have multiple causes, and they usually depend on sequencing protocol. As a result, estimating a realistic confidence score is critical for avoiding incorrect conclusions when analyzing RNA-seq data. We proposed the ‘positional bootstrap’, a method that computes confidence scores by simultaneously taking into account coverage and sequencing artifacts in a robust, general manner, that works even for unknown sources of error. It can also be readily combined with other bias-correcting methods to capture the effect of residual biases unaccounted for. In addition, we proposed novel methods for visualizing sequencing artifacts and comparing estimation methods across datasets. In these comparisons, the positional bootstrap outperformed the standard method used in the community by a wide margin, especially for the most confident estimates. These comparisons include analyses of alternative splicing, tissue-dependent splicing differences and tissue-dependent junction expression differences. Moreover, the positional bootstrap idea can potentially be applied to many other problems that make use of short reads, such as DNA sequencing, other RNA-seq tasks, CLIP-Seq and ChIP-Seq. The only requirement is that the estimator depends in the same way on some measurements obtained for a set of genomic positions. Bootstrapping these positions and combining the estimates of bootstrap samples produces a much more realistic confidence measure, by taking into account the empirical discrepancy between the positions due to sequencing artifacts. These sequencing artifacts can go undetected only if they happen to be the same across all genomic positions being analyzed, which is unlikely the case for many types of sequencing artifacts. Given the simplicity, rational and performance gain achieved by the positional bootstrap, we believe it will have wide applicability in the field.

## Acknowledgement

We thank Babak Alipanahi, Boyko Kakaradov and other members of Frey’s group for helpful discussions. This work was supported by CIHR, NSERC, John C. Polanyi and Ontario Genomics Institute (OGI) funding to BJF.

